# A family of RRM-1 RNA binding proteins enables cold adaptation and environmental resilience in *Bacteroides*

**DOI:** 10.64898/2026.07.07.737135

**Authors:** Hyelan Lee, Anubhav Basu, Carin K. Vanderpool

## Abstract

Bacteria use post-transcriptional regulatory mechanisms to rapidly adjust gene expression during environmental change. In the gut-associated genus *Bacteroides,* these mechanisms remain poorly defined as these organisms lack canonical RNA chaperones like Hfq and CsrA that coordinate post-transcriptional stress responses in many well-studied model bacteria. Most *Bacteroides* possess conserved RNA recognition motif-1 (RRM-1) domain-containing RNA-binding proteins (more common in eukaryotes than bacteria) that have been proposed to act as global RNA chaperones. Here, we show that these RNA binding proteins (RBPs) are central to cold stress adaptation. Simultaneous deletion of all *rbp* genes produces a cold-sensitive growth defect across multiple *Bacteroides* species, while single deletions do not, revealing conserved functional redundancy. RBP transcripts and proteins accumulate rapidly after temperature downshift, and loss of RBPs extensively reprograms the transcriptome. Cold sensitivity of *Bacteroides rbp* mutants is not caused by defects in ribosome assembly or rRNA maturation. Instead, we find that in *Bacteroides thetaiotaomicron,* RBPs act together with BT1884, the sole canonical cold shock protein possessed by this organism. The combined loss of RBPs and BT1884 produces a synthetic severe cold sensitivity phenotype, defining two functionally redundant cold stress systems belonging to unrelated protein families. Strains lacking RBPs show reduced survival under simultaneous cold and oxygen stress, the conditions *Bacteroides* cells are expected to encounter during host-to-host transmission. Together, these findings establish RRM-1 RBPs as non-canonical cold shock proteins that enable cold adaptation and environmental survival in *Bacteroides* and suggest how these organisms withstand the stresses of transmission between hosts.

**IMPORTANCE:** *Bacteroides* species are among the most abundant and stable members of the human gut microbiome, and they are also among the most readily transmitted between people. Reaching a new host requires surviving conditions outside the gut, including cold and oxygen exposure, yet how these bacteria withstand such stress is not well understood. Most bacteria manage stress using a well-defined set of RNA-binding proteins, but *Bacteroides* lack these canonical factors. We show that *Bacteroides* instead rely on a different family of RNA-binding proteins, more typical of eukaryotes than bacteria, to survive cold stress, and that these proteins promote survival under the conditions encountered during transmission. This work identifies a molecular system that allows an abundant and ecologically successful gut bacterium to endure the environmental challenges of moving between hosts.

## INTRODUCTION

It is well established that the gut microbiota exerts an extensive influence on host physiology, with reciprocal effects by the host and the environment on the structure and function of these microbial communities (1–3). Accordingly, substantial scientific interest has been directed toward defining the ecological and molecular determinants that guide colonization dynamics and microbial community assembly. Successful gut colonizers often possess specialized systems that confer fitness advantages and support their persistence in the host ecosystem (4–7). For example, *Bacteroides* species, among the most abundant and stable members of the human gut microbiota (8), have been shown to deploy many adaptive strategies, such as systems for use of complex diet- and host-derived polysaccharides (6, 9), stress response systems (10–13), and phase variable surface structures (7, 14, 15), that collectively support their colonization and maintenance in the gut environment. Recent studies demonstrate that *Bacteroides* can be horizontally transmitted between adult hosts (16, 17). During transmission, cells will be exposed to environmental stresses such as oxygen and temperature shifts. Thus, factors that enable cells to survive these environmental stresses during transmission likely provide a fitness advantage. While many studies have defined the intricate gene regulatory networks that control carbohydrate use (6, 9, 18), stress responses (10–13), and capsular phase variation (7, 14, 15) in *Bacteroides*, most prior work has focused on transcriptional control. However, the rapid environmental changes encountered by gut bacteria, together with the widespread use of post-transcriptional regulatory mechanisms across diverse bacteria (19, 20) underscore the importance of understanding post-transcriptional regulatory processes that enable *Bacteroides* to achieve rapid, energy-efficient physiological responses. Several studies have recently revealed the presence of hundreds of putative small regulatory RNAs (sRNAs) in *Bacteroides*, a subset of which have been functionally characterized (21–24). Yet, despite the growing body of evidence for post-transcriptional regulation in *Bacteroides,* canonical RNA chaperones including the widely conserved global regulators such as Hfq (25), ProQ (26), and CsrA (27), have not been identified. Previously, we reported the identification of RNA recognition motif-1 (RRM-1) domain-containing RNA binding proteins (RBPs) and found evidence supporting their potential roles as RNA chaperones (28). These RBPs are widely conserved in Bacteroidota and a recent study by Rüttiger et al., (24) provided key functional evidence supporting the role of RBPs as global RNA chaperones. Using CLIP-seq, they demonstrated that RbpB directly associates with over 170 mRNAs and 50 sRNAs. Furthermore, deletion of *rbpB* resulted in a colonization defect in male mice on a low fiber diet (24), highlighting the physiological relevance of RbpB in vivo.

RBPs may also perform other functions in post-transcriptional regulation, including roles typically associated with cold shock proteins (CSPs). During cold stress, bacteria face a global increase in the stability of RNA secondary structures that inhibits transcription elongation and translation; CSPs counteract this by binding single-stranded RNA and melting inhibitory structures to maintain efficient gene expression (29, 30). Canonical CSPs, which are defined by the presence of cold-shock domains (CSDs), are highly conserved across all domains of life (31). While classical CSPs such as CspA are archetypal cold-inducible proteins, other members of the CSP family in bacteria exhibit broader functions in general RNA metabolism, growth-phase regulation, and diverse stress responses beyond temperature downshift (29, 32–34). While many enteric bacteria encode multiple CSPs that contribute to cold adaptation (35–37), *Bacteroides thetaiotaomicron* encodes only a single annotated CSP: BT1884, which is a CspC/E homolog. In *E. coli*, CspC and CspE are expressed at 37°C and have been implicated in broader RNA-regulatory and stress-response functions, including regulation of *rpoS* and *uspA* expression, rather than functioning solely as cold-induced cold-shock proteins (38, 39). Thus*, B. thetaiotaomicron* may lack a canonical cold-inducible CSP repertoire, raising the question of how this organism manages cold stress with such a minimal canonical CSP repertoire. This raises the possibility that alternative RNA chaperones may compensate for cold stress-responsive functions in *Bacteroides*, and that RRM-1 RBPs may be among the proteins that fill this role.

RRM-1 domain-containing proteins are one of the most prevalent types of RNA-binding proteins in eukaryotes, and participate in many post-transcriptional activities including pre-mRNA splicing, mRNA export, mRNA stability and decay, and translational control (40). In bacteria, RRM-1 domain proteins exhibit a sparse yet broad distribution, occurring across multiple phyla rather than being restricted to a specific lineage (28, 41–45). The limited studies on bacterial RRM-1 proteins have implicated them in roles in stress and regulatory responses through post-transcriptional RNA-mediated regulation (28, 42–44). In cyanobacteria, where RRM-1 proteins are particularly prevalent, individual RBPs mediate stress responses including cold acclimation, circadian regulation, and salt tolerance, collectively acting as specialized RNA chaperones for environmental adaptation (45–47). Similarly, in *Porphyromonas gingivalis,* a member of the phylum Bacteroidota that also lacks canonical RNA chaperones, an RRM-1 domain protein has been implicated in virulence factor regulation and intracellular persistence (42). Together, these observations suggest that RRM-1 domain proteins can mediate diverse stress-responsive functions through post-transcriptional regulation, but whether RRM-1 RBPs in *Bacteroides* contribute to similar stress adaptation processes remains unknown.

Here, we identify a previously unrecognized role for RRM-1 domain-containing RBPs in *Bacteroides* cold adaptation. Reasoning that *Bacteroides* species have a minimal CSP repertoire yet must mount rapid post-transcriptional responses to environmental stress, we asked whether RRM-1 RBPs fulfill the cold stress adaptation functions typically carried out by canonical CSPs. We find that simultaneous loss of all RBPs produces a pronounced cold-sensitive phenotype conserved across multiple *Bacteroides* species, and that *rbp* transcription and RBP protein production are induced upon temperature downshift. Loss of RBPs reshapes the transcriptome far more extensively at low temperature than at 37°C, yet does not impair ribosome assembly or rRNA maturation, indicating that RBPs support cold adaptation through a mechanism distinct from the ribosome biogenesis defects commonly associated with cold sensitivity. We further show that RBPs function in coordination with BT1884, the sole canonical CSP encoded by *B. thetaiotaomicron,* revealing that *Bacteroides* deploys two functionally redundant cold stress systems from unrelated protein families. Finally, we find that RBPs promote *Bacteroides* survival under the combined cold and oxygen stress that *Bacteroides* cells encounter outside the host. Together, these findings establish RRM-1 RBPs as non-canonical cold shock proteins that enable cold adaptation and environmental survival in *Bacteroides*.

## RESULTS

### Loss of RBPs results in a cold sensitivity across *Bacteroides* species

The RRM-1 domain containing RBPs are broadly distributed across *Bacteroides*, with most species encoding multiple *rbp* copies per genome (28). Previously, we showed that *B. thetaiomiocron* encodes three RRM-1 domain-containing RBPs: RbpA (BT0784), RbpB (BT1887), and RbpC (BT3840) (28). Given the possibility that *Bacteroides* RBPs contribute to cold shock responses, we first examined whether their loss leads to a cold-sensitive phenotype. We compared the growth of *B. thetaiotaomicron* wild-type (WT) and *rbp* deletion strains at 37°C (optimal growth temperature) and 21°C (cold stress) and found that while strains lacking a single RBP (Δ*rbpA*, Δ*rbpB*, Δ*rbpC*) showed no discernible growth defects at either 37 or 21°C compared to the WT strain, the triple deletion strain (Δ*rbpABC)* exhibited markedly impaired growth at 21°C, but not at 37°C, indicative of a cold-sensitive phenotype (Fig. 1A). Complementation of the Δ*rbpABC* strain with a single chromosomal copy of any individual *rbp* restored growth at 21°C, confirming that the cold-sensitive phenotype is specific to loss of RBP function and that each RBP is individually sufficient to support cold tolerance (Fig. S1).

**Figure 1.**
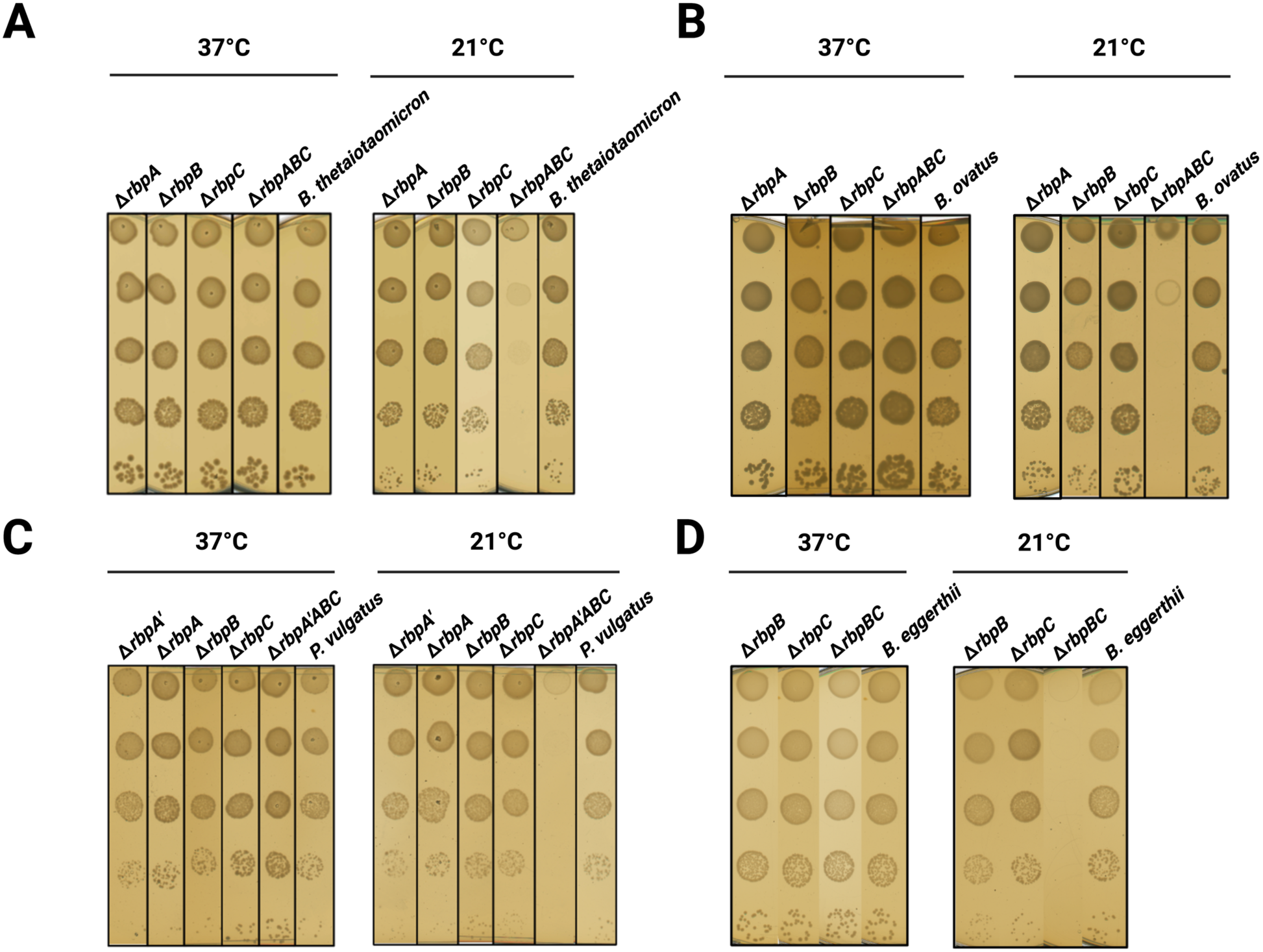
Cold-sensitive phenotype of RBP deletion mutants across *Bacteroides* species. Wild-type strains and mutants carrying either single *rbp* deletions or deletions of all *rbp* homologs were analyzed in (A) *B. thetaiotaomicron*, (B) *B. ovatus*, (C) *P. vulgatus,* and (D) *B. eggerthii.* Strains were grown anaerobically in TYG medium at 37°C to exponential phase. Cultures were serially diluted 10-fold, and 10 µl of each dilution was spotted onto BHI-S agar plates. Plates were incubated anaerobically at 37°C or 21°C for 2 days and 5 days, respectively, as indicated. Representative results from three biological replicates are shown.

To determine whether the cold-sensitive phenotype associated with RBP loss is conserved across the genus, we generated strains lacking individual or all *rbp* genes in several *Bacteroides* species with different *rbp* gene copy numbers. *B. thetaiotaomicron* and *Bacteroides ovatus* each possess three copies (*rbpA, rbpB*, and *rbpC*), the most prevalent copy number in *Bacteroides*. *Bacteroides eggerthii* possesses two copies (*rbpB* and *rbpC*) whereas *Phocaeicola vulgatus* (previously *Bacteroides vulgatus*) possesses four copies (*rbpA*, *rbpA’*, *rbpB*, and *rbpC*) (28). In all four species, strains lacking any single RBP grew comparably to the corresponding WT at both 37°C and 21°C, while strains lacking all *rbp* copies exhibited impaired growth specifically at 21°C (Fig. 1B, C, D). These results demonstrate that a cold-sensitive phenotype upon complete loss of RBPs is conserved across *Bacteroides* species encoding different numbers of *rbp* genes, establishing functional redundancy among RBPs as a conserved feature of the genus.

### RNA seq reveals extensive RBP-dependent transcriptome changes during cold stress

To investigate the molecular basis of the cold-sensitive phenotype observed in the Δ*rbpABC* mutant, we performed RNA-seq on *B. thetaiotaomicron* WT and Δ*rbpABC* strains grown at either 37°C or 21°C. Principal component analysis (PCA) revealed that temperature was the dominant factor influencing global transcription patterns, explaining 70% of the variance in the data (Fig. 2A). Consistent with this, a significant number of genes were temperature-responsive in both strains: 41% of genes (2024/4911) in WT and 49% (2402/4911) in Δ*rbpABC* were differentially expressed between 37°C and 21°C (absolute log2 fold change >1, adjusted p-value <0.05; Fig. S2A; Table S1 and S2). The larger fraction of temperature-responsive genes in the Δ*rbpABC* strain suggests that RBPs normally buffer a portion of the transcriptional response to cold.

**Figure 2.**
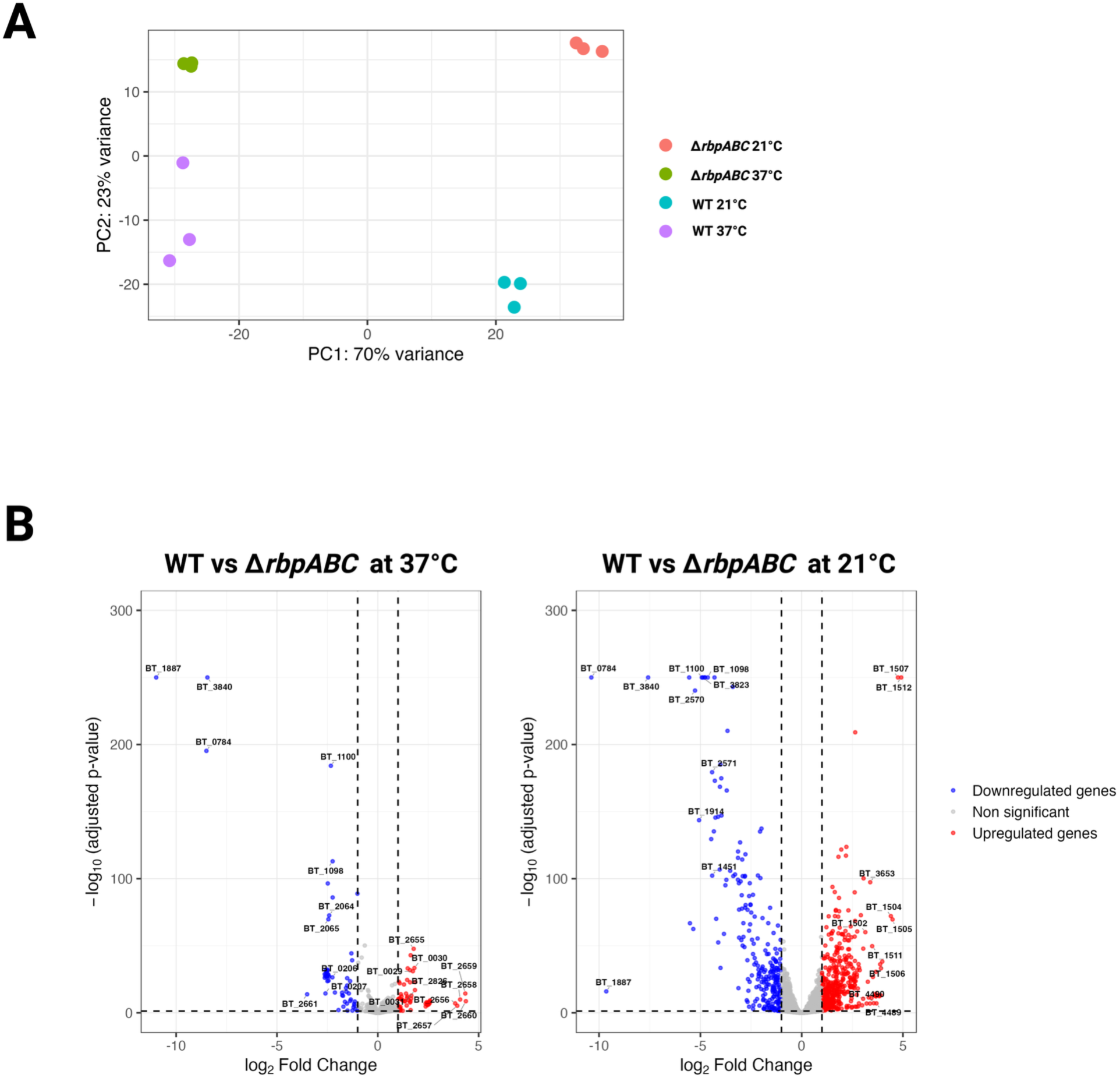
RNA-seq reveals extensive RBP-dependent transcriptomic changes specifically at cold temperature. (A) Principal component analysis (PCA) plot showing clustering of RNA-seq samples based on gene expression in WT and Δ*rbpABC* strains at 37°C and 21°C. (B) Volcano plots showing upregulated genes (red) and downregulated genes (blue) in the Δ*rbpABC* strain compared to WT at 37°C and 21°C. For visualization, adjusted *P* values less than 10^−250^ were set to 10^−250^. The top 10 upregulated and top 10 downregulated genes, ranked by log2 fold change, are labeled. Genes located within known capsular polysaccharide loci were excluded from labeling because these loci are subject to sporadic induction by invertible promoters.

To examine the specific impact of RBPs on gene regulation, we compared transcript levels between WT and Δ*rbpABC* strains at each temperature. At 37°C, 128 genes were differentially expressed between the two strains (72 upregulated, 56 downregulated). In contrast, at 21°C, 735 genes were differentially expressed (472 upregulated, 263 downregulated), 654 of which were uniquely altered at cold temperature (Fig. 2B; Table S3 and S4). This increase in RBP-dependent transcriptome changes at 21°C compared to 37°C demonstrates that RBPs exert a substantially greater regulatory impact under cold conditions, consistent with their roles as cold-responsive proteins.

To identify the biological processes most affected by RBP loss under cold conditions, we performed over-representation analysis (ORA) and gene set enrichment analysis (GSEA). At 37°C, no pathways were significantly enriched by GSEA, and ORA identified only genes in annotated capsular polysaccharide loci (Fig. S2B; Table S5 and S6), suggesting that RBPs have limited impact on major metabolic pathways at optimal growth temperature. At 21°C, however, multiple pathways were significantly downregulated in the Δ*rbpABC* mutant, including ATP metabolic processes, amino acid metabolism, aminoacyl-tRNA biosynthesis, and the pentose phosphate pathway (Fig. S2B; Table S5 and S6). Consistent with these pathway-level changes, genes encoding the F-type ATP synthase complex (*BT0711-BT0719*), glycolytic enzymes, and components of the pentose phosphate pathway were among the most strongly downregulated. These changes point to a coordinated reduction in energy-generating pathways in the Δ*rbpABC* mutant at 21°C. Because this strain also grows poorly at 21°C, we cannot distinguish whether the downregulation of energy metabolism contributes to the cold-sensitive growth defect or arises as a consequence of it.

### RBP transcripts and proteins are rapidly and strongly induced by cold stress

RNA-seq analysis revealed significant upregulation of all three *rbp* transcripts in WT cells grown at 21°C compared to 37°C, with approximately 11-fold induction of *rbpA* and 6-fold induction of *rbpB* and *rbpC* (Table S1). To determine whether this transcriptional induction is reflected at the protein level, we performed Western blot analysis on strains producing epitope-tagged RBPs. RbpA, RbpB, and RbpC protein levels were all markedly elevated at 21°C compared to 37°C, and RbpA and RbpB abundance were substantially reduced in stationary phase, indicating that RBP accumulation is coordinated with both temperature and growth state (Fig. 3A).

**Figure 3.**
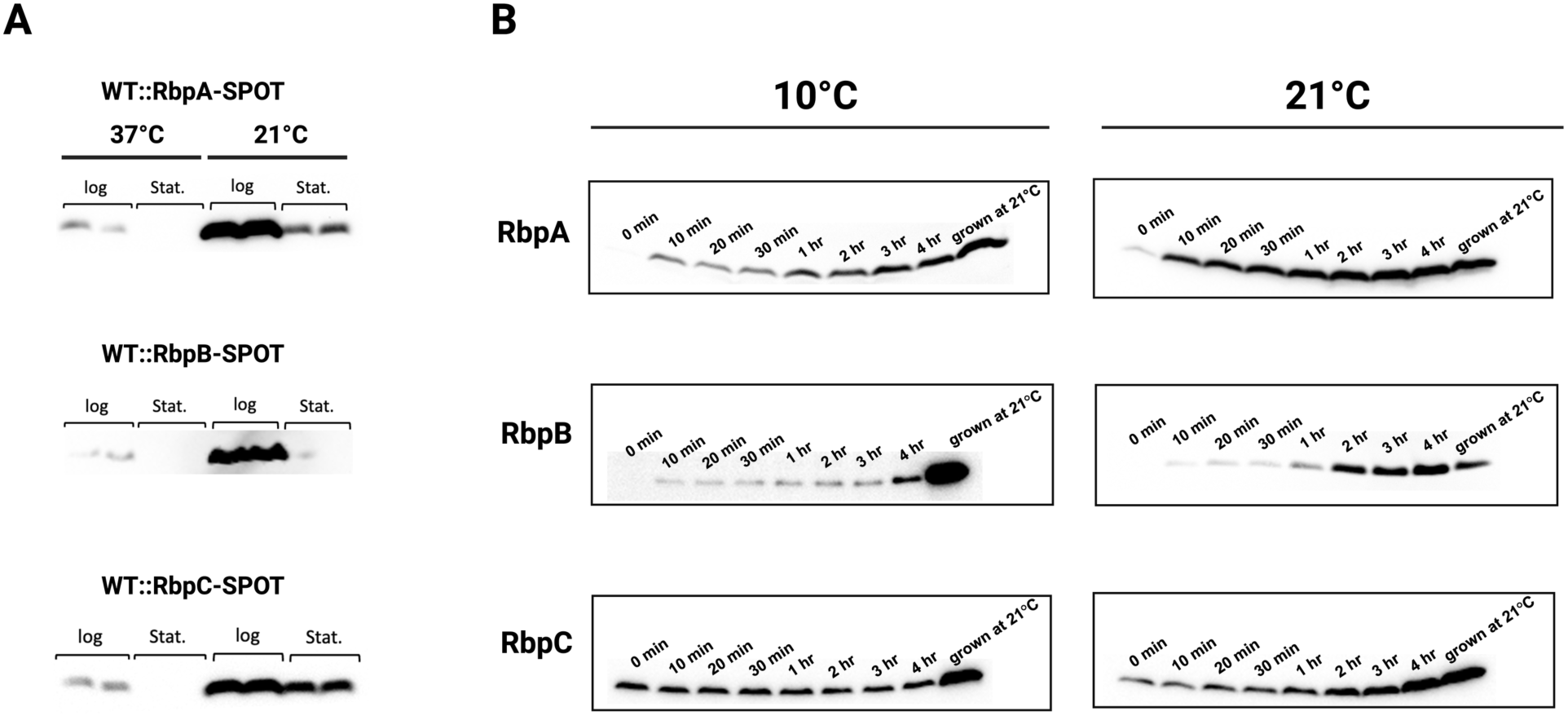
RBP protein levels are rapidly and strongly induced upon cold exposure. (A) Western blot analysis of RbpA-SPOT, RbpB-SPOT, and RbpC-SPOT levels in *B. thetaiotaomicron*. Strains carrying chromosomally SPOT-tagged *rbpA*, *rbpB*, or *rbpC* were grown anaerobically at 37°C or 21°C and harvested during exponential phase (log) or stationary phase (stat). Whole-cell lysates were separated by SDS-PAGE and analyzed by Western blotting with anti-SPOT antibody. (B) Time-course Western blot analysis of RbpA-SPOT, RbpB-SPOT, and RbpC-SPOT following temperature downshift. Strains were grown anaerobically at 37°C to exponential phase and then shifted to 21°C or 10°C. Cells were harvested at the indicated time points after temperature shift, and protein levels were evaluated by Western blotting using an anti-SPOT antibody.

To examine the kinetics of RBP induction, we monitored protein levels by Western blot following a temperature downshift from 37°C to either 21°C or 10°C. RbpA accumulated rapidly under both conditions, with increased levels detectable within 10 minutes of the temperature shift and continuing to accumulate thereafter; by 4 hours, RbpA levels were comparable to those in cells grown to exponential phase at 21°C (Fig. 3B, top panels). RbpB showed a similar pattern, rising from marginally detectable at 37°C (0 minutes) to clearly detectable within 10 minutes, though accumulation was slower and reached lower levels after a shift to 10°C than to 21°C (Fig. 3B, middle panels). RbpC behaved differently from RbpA and RbpB: it was present at higher basal levels at 37°C (0 minutes), showed no noticeable induction after a shift to 10°C, and accumulated more slowly at 21°C than either RbpA or RbpB (Fig. 3B, bottom panels).

While the kinetics of cold-inducibility differs between the three RBPs (Fig. 3B), it is clear that all three achieve higher steady-state levels at low temperature (Fig. 3A). These data, combined with the pronounced cold-sensitive phenotype of strains lacking RBPs (Fig. 1), strongly support a critical role for these proteins in cold adaptation. The differences in induction kinetics and temperature responsiveness suggest that each RBP is subject to distinct regulatory inputs, reminiscent of the functional heterogeneity seen among canonical CSP family members (48).

### RBPs are unlikely to play a role in ribosome assembly or maturation

Cold-sensitive phenotypes in bacteria frequently arise from defects in ribosome biogenesis (49–51), and RNA chaperones including Hfq have been shown to contribute to ribosome maturation (52). We tested whether impaired ribosome biogenesis might underlie the observed cold-sensitive phenotype of the *B. thetaiotaomicron* Δ*rbpABC* mutant and found four independent lines of evidence that argue against this model. First, sucrose gradient sedimentation analysis revealed that the proportions of ribosomal subunits and 70S monosomes in the Δ*rbpABC* strain were indistinguishable from WT at both 37°C (not shown) and 21°C (Fig. 4), indicating that overall ribosome assembly is unaffected. Second, total RNA from Δ*rbpABC* cells showed no accumulation of 17S rRNA, suggesting that rRNA maturation is intact (Fig. S3A). Third, our RNA-seq data revealed no differential abundance of ribosomal protein transcripts in the Δ*rbpABC* background, in contrast to what has been reported for *hfq* mutants (52) (Table S3 and S4). Finally, overexpression of *B. thetaiotaomicron* genes encoding homologs of canonical ribosome assembly factors known to suppress temperature-sensitive ribosome biogenesis defects, including *rsgA*, *rbfA*, *dnaK*, *der*, *rsmA*, *era*, *rimP*, and *rimM*, failed to rescue the cold-sensitive phenotype of the Δ*rbpABC* strain (Fig. S3B) (50, 51, 53–57). Together, these results indicate that the cold sensitivity of the Δ*rbpABC* strain arises through a mechanism distinct from impaired ribosome biogenesis.

**Figure 4.**
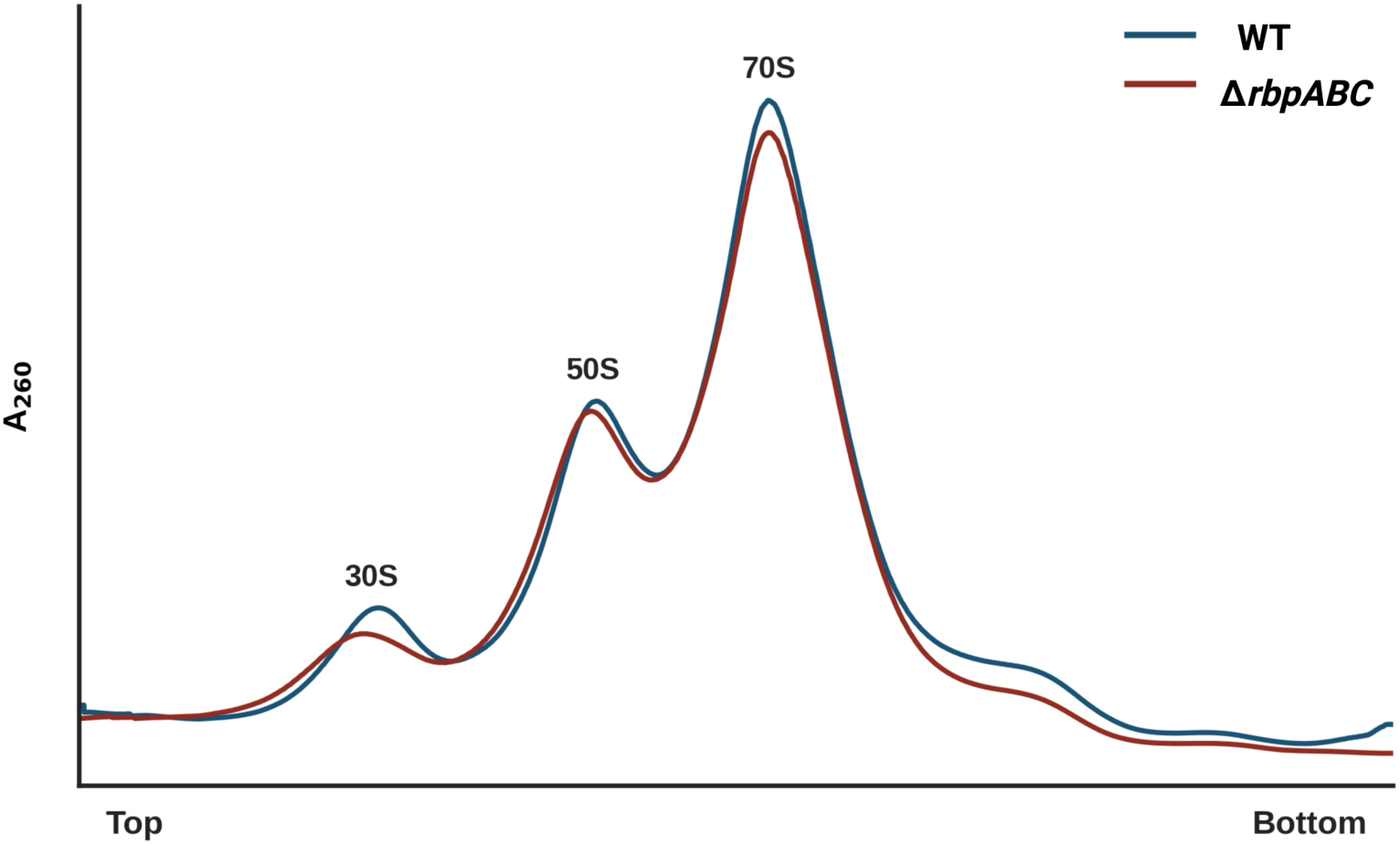
Loss of RbpA, RbpB, and RbpC does not substantially alter ribosome sedimentation profiles. Shown are sucrose gradient sedimentation profiles of ribosomes isolated from WT and Δ*rbpABC* strains grown at 21°C. Absorbance at 260 nm was monitored across the gradient. The positions of the 30S and 50S ribosomal subunits and 70S monosomes are indicated. WT and Δ*rbpABC* strains showed similar 30S, 50S, and 70S peak patterns.

### Loss of BT1884 exacerbates cold sensitivity in the absence of RBPs

Having established that RBPs are required for cold tolerance and are strongly induced at low temperature, we next asked whether *B. thetaiotaomicron* encodes additional factors that contribute to cold adaptation. BT1884, the sole CSP encoded by *B. thetaiotaomicron* and a putative homolog of *E. coli* CspC and CspE, was a natural candidate. Notably, *BT1884* is co-transcribed with *BT1885* (a DEAD/DEAH box helicase), *BT1886* (a nitronate monooxygenase), and *BT1887* (RbpB) as a single operon (28), suggesting a potential functional connection between BT1884 and RBPs.

We constructed Δ*BT1884* and Δ*BT1884-1887* operon deletion strains in both WT and Δ*rbpABC* backgrounds and compared their growth at 37°C and 21°C. At 37°C, none of the strains displayed a growth defect (Fig. 5, left panel). At 21°C, deletion of *BT1884* or the entire *BT1884-1887* operon in an otherwise WT background had no detectable effect on growth. However, in the Δ*rbpABC* background, additional deletion of *BT1884* markedly exacerbated the cold-sensitive phenotype, producing a synthetic cold-sensitivity not observed at 37°C (Fig. 5, right panel). Deletion of the full *BT1884-1887* operon did not worsen the defect beyond that of Δ*rbpABC* Δ*BT1884*, indicating that BT1884 is the primary contributor to cold adaptation within the operon and that BT1885 and BT1886 play minimal roles under these conditions.

**Figure 5.**
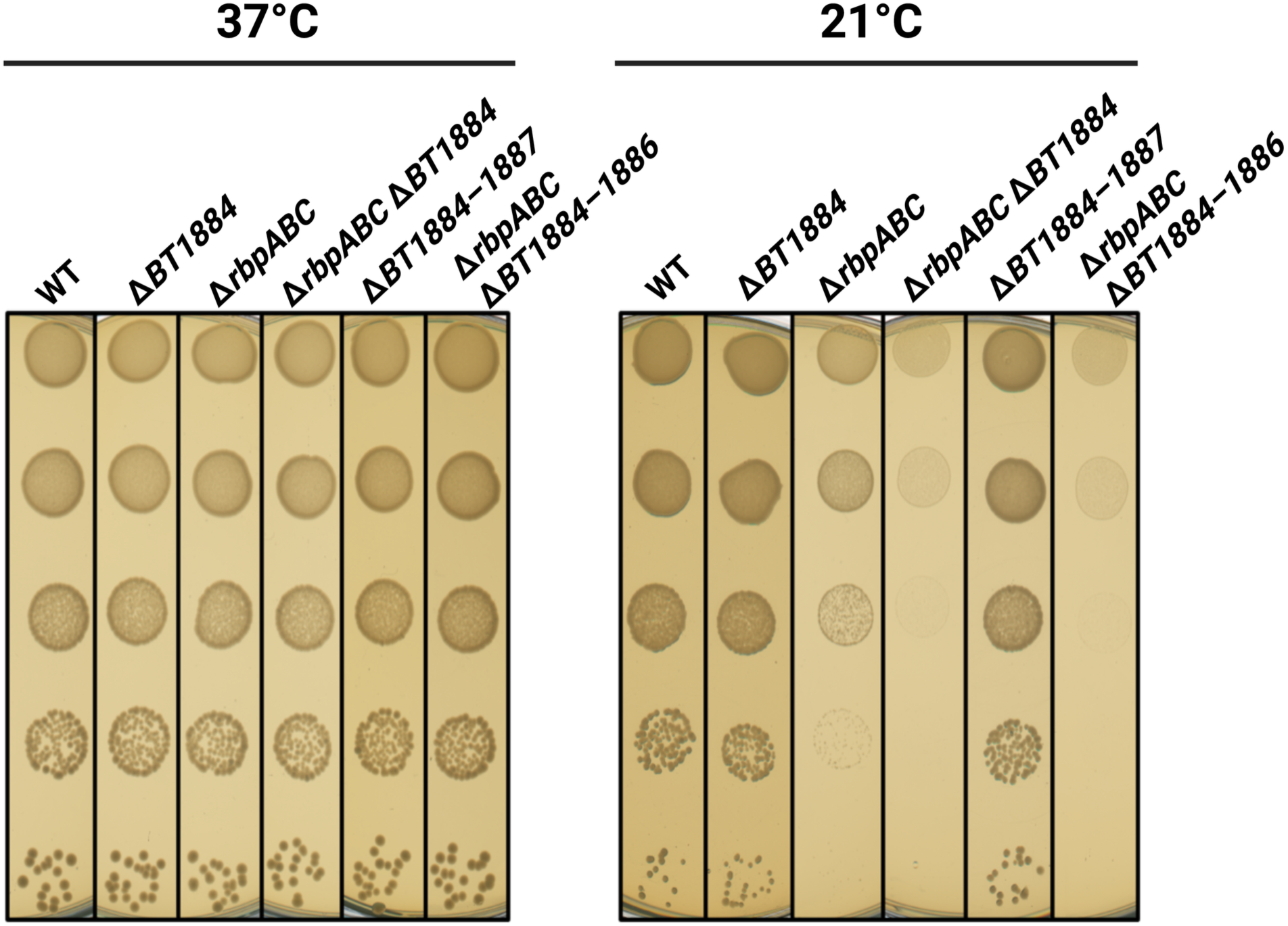
*BT1884* deletion exacerbates the cold-sensitive phenotype of Δ*rbpABC*. *B. thetaiotaomicron* WT, Δ*BT1884*, Δ*rbpABC*, Δ*rbpABC* Δ*BT1884*, Δ*BT1884*–*1887*, and Δ*rbpABC* Δ*BT1884*–*1886* strains were grown anaerobically in TYG medium at 37°C to exponential phase. Cultures were serially diluted 10-fold, and 10 µl of each dilution was spotted onto BHI-S agar plates. Plates were incubated anaerobically at 37°C or 21°C for 2 days and 5 days, respectively, as indicated. *BT1887* corresponds to *rbpB* and is already deleted in the Δ*rbpABC* background. Representative results from three biological replicates are shown.

These results demonstrate that BT1884 and RBPs act in a partially redundant manner to support cold tolerance in *B. thetaiotaomicron*, revealing that this organism relies on two functionally distinct cold stress systems, the RRM-1 RBPs and its sole canonical CSP, that together ensure robust adaptation to temperature downshift.

### RBPs regulate their own expression through redundant feedback involving both transcriptional and post-transcriptional control

Expression of canonical CSPs in other bacteria is governed by both transcriptional regulation and post-transcriptional mechanisms operating through their long 5’ UTRs, and *Bacteroides* RBP expression may similarly be subject to multi-level control. In *E. coli*, CspA autoregulates its own abundance through negative feedback, and loss of individual CSPs can trigger compensatory induction of other family members (37, 58, 59). The *rbp* transcripts in *B. thetaiotaomicron* harbor 5’ UTRs of 135 nt for *rbpA*, 122 nt for *rbpB*, and 155 nt for *rbpC* (21), comparable in length to the 159 nt 5’ UTR of *E. coli cspA* (60). To test for auto- and cross-regulation mediated by *rbp* 5’ UTRs, we constructed nanoluciferase (NanoLuc) reporter fusions integrated into a neutral chromosomal locus in WT and Δ*rbp* backgrounds to monitor *rbp* transcription and translation (Fig. 6A). “Native” reporters contained the native *rbp* promoter, 5ʹUTR, and the coding sequence for the first 10 amino acids fused to NanoLuc and reporter activity will reflect combined contributions of transcriptional and translational regulation. Transcriptional fusions contained only the native *rbp* promoter fused to NanoLuc to measure solely transcriptional regulation, whereas translational fusions contained the *rpoD* promoter, the native *rbp* 5ʹUTR, and the coding sequence for the first 10 amino acids fused to NanoLuc to isolate post-transcriptional regulatory processes mediated by the 5ʹUTR. At 37°C, native *rbp* reporter activity was largely similar between WT and Δ*rbp* strain backgrounds, although modest increases were observed for all three native fusions in the Δ*rbpABC* background and for the *rbpA* native fusion in the Δ*rbpC* background (Fig. 6B). This increase in the Δ*rbpABC* background disappeared upon complementation with RbpC. At 21°C, native fusion activity was similar between the WT strain and strains with a single *rbp* gene deletion, and substantially increased in the Δ*rbpABC* background, with approximately 2.2-fold, 2.5-fold, and 3-fold higher activity for *rbpA*, *rbpB*, and *rbpC*, respectively, compared with WT (Fig. 6C). For the *rbpC* native fusion, the increased reporter activity in the Δ*rbpABC* background was reversed by complementation with a single chromosomal copy of *rbpC* (Fig. 6C). These data suggest that *rbp* expression may be subject to negative auto- and cross-regulation.

**Figure 6.**
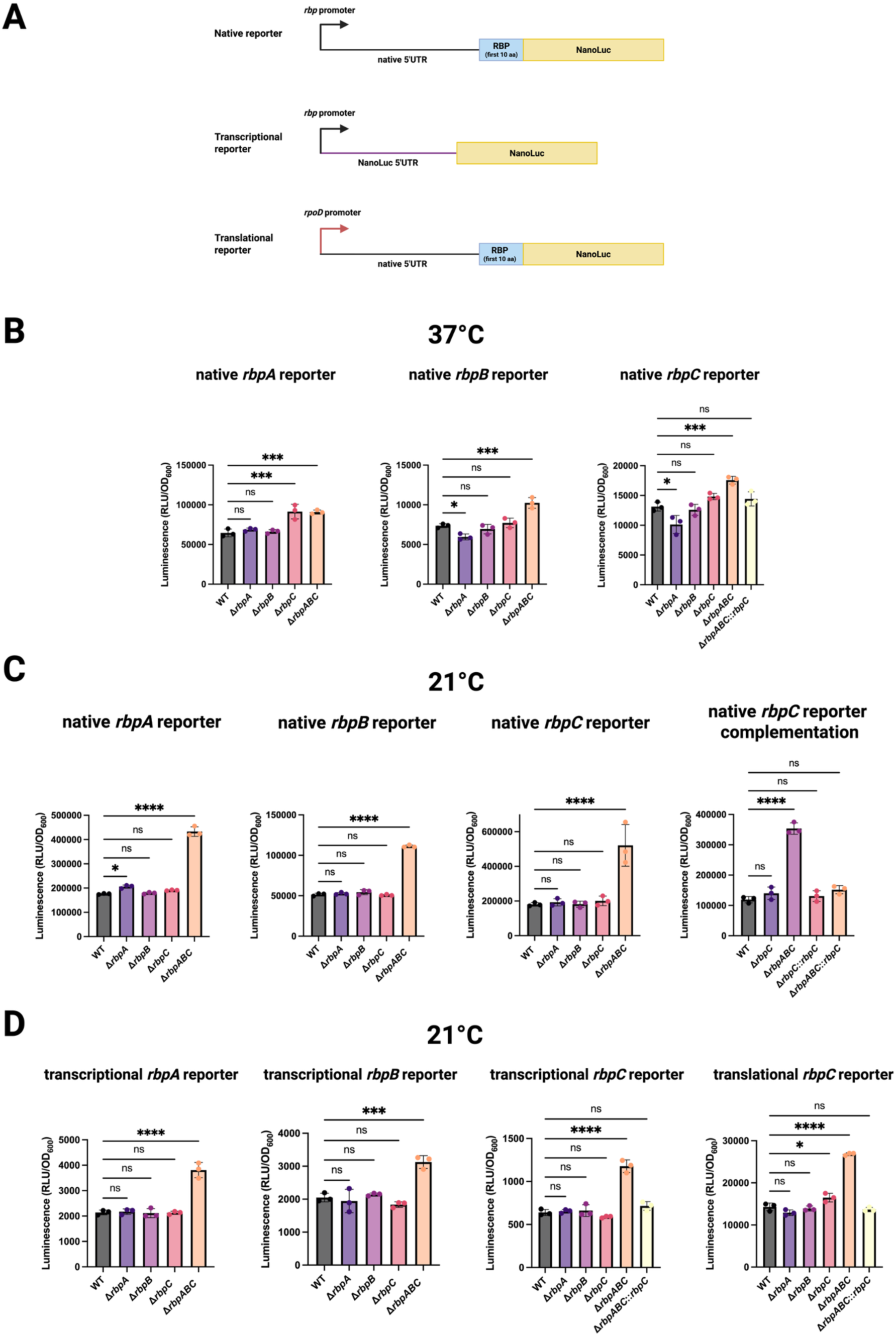
*rbp* expression is modulated by redundant regulatory mechanisms involving both transcriptional and post-transcriptional regulation. (A) Schematic of the reporter fusion constructs used in this study. Native fusions contain the native *rbp* promoter, 5′ UTR, and the coding sequence for the first 10 amino acids fused to NanoLuc. Transcriptional fusions contain the native *rbp* promoter fused to NanoLuc. Translational fusions contain the *rpoD* promoter, the native *rbp* 5′ UTR, and the coding sequence for the first 10 amino acids fused to NanoLuc. (B) NanoLuc reporter activity from native *rbpA*, *rbpB*, and *rbpC* fusions in WT, *rbp* deletion mutants, and complemented strains grown anaerobically at 37°C. (C) NanoLuc reporter activity from native *rbpA*, *rbpB*, and *rbpC* fusions in WT, *rbp* deletion mutants, and complemented strains grown anaerobically at 21°C. (D) NanoLuc reporter activity from transcriptional *rbpA*, transcriptional *rbpB*, transcriptional *rbpC*, and translational *rbpC* fusions in WT, *rbp* deletion mutants, and complemented strains grown anaerobically at 21°C. Reporter activity was measured during exponential phase and expressed as relative luminescence units normalized to OD600. Error bars represent the standard deviations of three biological replicates, and statistical significance was determined by ordinary one-way ANOVA followed by Dunnett’s multiple-comparisons test comparing each strain to WT. P values are indicated as follows: *P* < 0.05 (*), *P* < 0.01 (**), *P* < 0.001 (***), *P* < 0.0001 (****).

To determine whether auto- and cross-regulation occurs at the transcriptional or post-transcriptional level (or both), we measured activity of additional reporter constructs for each *rbp* gene (Fig. 6A). At 21°C, the *rbpA* and *rbpB* transcriptional fusion activities were similar between the WT strain and strains with a single *rbp* gene deletion and showed increased activity in the Δ*rbpABC* background. The magnitude of increased activity in the Δ*rbpABC* background was only 1.8 and 1.5-fold for *rbpA* and *rbpB* fusions, respectively, compared to 2.5 and 2.2-fold for the corresponding native fusions (Fig. 6D). A similar pattern was observed for *rbpC* where we compared activities of both transcriptional and translational fusions. There were no substantial differences in fusion activity between WT and single *rbp* deletion strains, and activity in the Δ*rbpABC* background was increased, but to a more modest degree compared with the native fusion (approximately 2-fold versus 3-fold; Fig. 6D). These results indicate that promoter-dependent transcriptional regulation contributes to RBP feedback control but does not fully account for the stronger induction observed with the native fusions. The induction of the *rbpC* translational fusion further suggests that the native 5ʹUTR also contributes to post-transcriptional regulation. Together, these findings support a model in which redundant feedback regulation of *rbp* expression under cold conditions involves both transcriptional and post-transcriptional control. The fact that de-repression is observed only upon loss of all three RBPs indicates that each RBP contributes to maintaining repression of itself and other RBPs.

### BT1884 modulates RBP expression under cold stress

Given that loss of BT1884 synthetically enhanced the cold-sensitive phenotype of the Δ*rbpABC* strain, we next asked whether BT1884 also influences *rbp* expression. We introduced an *rbpC* native reporter fusion into WT, Δ*rbpABC*, Δ*BT1884*, and Δ*rbpABC* Δ*BT1884* strains and measured activity at 21°C. As previously shown, the *rbpC* native fusion exhibited an approximately 3-fold increase in activity in the Δ*rbpABC* strain relative to the WT strain (Fig. 7). Activity of the fusion in the Δ*BT1884* strain was similar to activity in the wild-type strain. In the Δ*rbpABC* Δ*BT1884* strain, fusion activity was intermediate: higher than in the WT and Δ*BT1884* strains, but lower than in the Δ*rbpABC* strain (Fig. 7). These data suggest that BT1884 contributes to the increased *rbpC* expression observed in the absence of RBPs.

**Figure 7.**
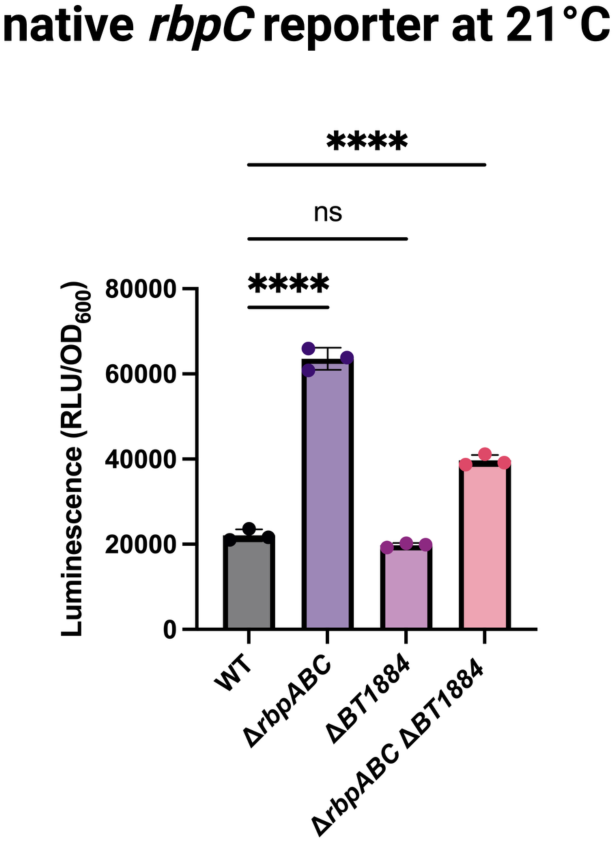
BT1884 modulates *rbp* expression under cold stress. A NanoLuc reporter containing the native *rbpC* promoter, 5′ UTR, and coding sequence for the first 10 amino acids was integrated at a neutral chromosomal locus in the indicated strains. Strains were grown anaerobically at 21°C. Reporter activity was measured during exponential phase and expressed as relative luminescence units normalized to OD600. Error bars represent the standard deviations of three biological replicates, and statistical significance was determined by ordinary one-way ANOVA followed by Dunnett’s multiple-comparisons test comparing each strain to WT. P values are indicated as follows: P < 0.05 (*), P < 0.01 (**), P < 0.001 (***), P < 0.0001 (****).

These results reveal a functional connection between BT1884 and RBP-mediated feedback regulation under cold conditions. Given that BT1884 is a CspC/E homolog and the sole canonical CSP in *B. thetaiotaomicron*, it may influence *rbp* expression through effects on RNA stability or translation efficiency analogous to those described for CSP family members in other bacteria (30, 61, 62). Together with the genetic interactions observed in the cold-sensitivity phenotypes, these findings suggest that BT1884 and RBPs operate within a shared regulatory network that collectively governs cold adaptation in *Bacteroides*.

### RBPs promote *Bacteroides* survival under cold and oxygen stress

During host-to-host transmission, *Bacteroides* cells are likely exposed to environments dissimilar from the gut and encounter various stresses such as cold and oxygen exposure. Given that RBPs mediate cold adaptation, we next tested whether RBPs influence the survival of *Bacteroides* under conditions that may resemble extra-host environments. WT, Δ*rbpABC*, Δ*BT1884*, and Δ*rbpABC* Δ*BT1884* strains were monitored for long-term viability. Strains were grown anaerobically to stationary phase at 37°C and then transferred to 37°C anaerobic, 21°C anaerobic, or 21°C aerobic conditions. Viable cell counts were measured at days 0, 6, 12, and 18 and expressed as a percentage of the starting CFU/ml for each biological replicate (Fig. 8). At 37°C, all four strains showed a comparable decline in viability, with no significant differences detected among strains at any time point. This suggests that loss of RBPs or BT1884 does not confer a survival defect at 37°C under anaerobic conditions. At 21°C, viability remained similar across strains through day 6. However, by day 12, the Δ*rbpABC* strain showed a statistically significant reduction in viability compared with WT (2.17 % survival for Δ*rbpABC* and 4.08 % survival for WT), and this difference persisted through day 18 (0.82 % survival for Δ*rbpABC* and 1.76 % survival for WT). The Δ*BT1884* single mutant also showed a modest but statistically significant reduction in viability at day 18 (1.24 % survival) compared with WT. The Δ*rbpABC* Δ*BT1884* mutant showed a reduction in viability similar to Δ*rbpABC* by day 18 (0.91 % survival). The most severe phenotypes were observed under combined cold and oxygen stress. Both Δ*rbpABC* and Δ*rbpABC* Δ*BT1884* displayed significantly reduced viability relative to WT by day 12 (0.0064 % survival for Δ*rbpABC*, 0.0059 % survival for Δ*rbpABC* Δ*BT1884*, and 0.025 % survival for WT). By day 18, survival differences were larger: WT survival was 11.7-fold higher than Δ*rbpABC*, and 33.6-fold higher than Δ*rbpABC* Δ*BT1884* (0.000087 % survival for Δ*rbpABC*, 0.00003 % survival for Δ*rbpABC* Δ*BT1884*, and 0.001 % survival for WT). In contrast, the Δ*BT1884* single mutant did not differ significantly from WT under combined cold and aerobic stress at any time point, indicating that *BT1884* is not independently required for survival under this condition. The substantially greater impairment of Δ*rbpABC* Δ*BT1884* compared with Δ*rbpABC* under 21°C aerobic conditions suggests that *BT1884* may provide additional protection against combined cold and aerobic stress in the absence of functional RBPs.

**Figure 8.**
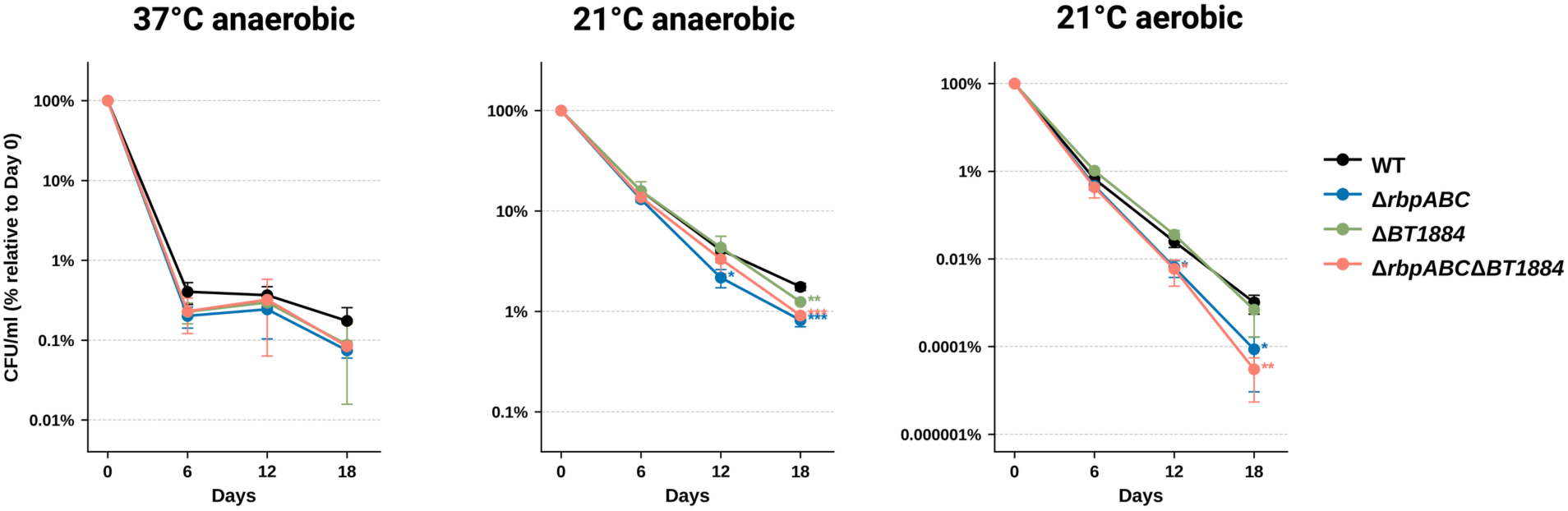
RBPs and the cold shock protein homolog *BT1884* contribute to *Bacteroides thetaiotaomicron* survival under cold and aerobic stress. *B. thetaiotaomicron* WT, Δ*rbpABC*, Δ*BT1884*, and Δ*rbpABC* Δ*BT1884* strains were grown to stationary phase at 37°C and then transferred to the indicated conditions. Viable cell counts (CFU/ml) were measured at days 0, 6, 12, and 18. Data are expressed as the percentage of CFU/ml relative to the day 0 count for each biological replicate. Points and error bars represent means ± standard deviations of three independent biological replicates, except for Δ*BT1884* and Δ*rbpABC* Δ*BT1884* at day 12, where *n* = 2. Statistical comparisons between each mutant strain and WT at each time point were performed using one-way ANOVA with Dunnett’s multiple-comparisons test applied to log_10_-transformed viability values. *P* values are indicated as follows: *P* < 0.05 (*), *P* < 0.01 (**), *P* < 0.001 (***).

Together, these results indicate that RBPs are important determinants of *B. thetaiotaomicron* survival during cold stress, particularly when cold stress is combined with oxygen exposure. Although *BT1884* was not independently required for survival under combined cold and oxygen stress, the enhanced defect of the Δ*rbpABC* Δ*BT1884* mutant suggests that *BT1884* may provide additional protection when RBP function is absent. These findings support the hypothesis that RBPs, and potentially broader cold stress response pathways, may influence fitness during transmission events that expose *Bacteroides* cells to environments outside the host.

## DISCUSSION

This study establishes RRM-1 RBPs as non-canonical cold shock proteins in *Bacteroides*, revealing a functionally redundant regulatory system that is rapidly induced by temperature downshift and conserved across *Bacteroides* species encoding different *rbp* copy numbers. The cold sensitive phenotype conferred by deletion of all *rbp* copies was conserved across four species spanning two genera, *Bacteroides* and *Phocaeicola*, and across strains encoding two, three, or four *rbp* copies. This breadth of conservation, across both phylogenetic distance and gene copy number, indicates that redundant RBP function is an ancient and maintained feature of this clade rather than a property of any single species, and argues that the cold adaptation role we describe is likely to operate broadly across the Bacteroidota. The finding that complete loss of RBPs is required to produce a cold-sensitive phenotype and that a single RBP is sufficient to restore cold tolerance points to a robust and buffered regulatory architecture. The genetic relationship between RBPs and BT1884, the sole canonical CSP encoded by *B. thetaiotaomicron* is notable. The combined loss of RBPs and BT1884 produces a severe synthetic cold sensitive phenotype, and BT1884 contributes to the upregulation of RBP expression observed in the Δ*rbpABC* background under cold conditions. Together, these findings suggest that *Bacteroides* has compensated for its minimal canonical CSP repertoire by deploying functionally distinct but partially redundant cold stress factors from different protein families, a solution that appears to be broadly conserved across the genus.

The parallels between RRM-1 RBPs and canonical CSPs are striking and provide many avenues for future investigation. Like CspA in *E. coli* (30), the RBPs harbor long 5’ UTRs of comparable length to the *cspA* 5’ UTR and are subject to feedback regulation that restricts their own expression under cold conditions. Our reporter assays demonstrate that this feedback operates at both transcriptional and post-transcriptional levels and requires the combined activity of all three RBPs to maintain repression. This redundant regulatory architecture mirrors the functional redundancy observed in the cold-sensitive growth phenotype. However, the three RBPs are not uniform in their cold responsiveness. RbpA and RbpB are rapidly and strongly induced at both the transcript and protein levels following temperature downshift, whereas RbpC is present at higher basal levels and is induced more modestly. Heterogeneity in cold-inducibility is also a feature of canonical CSP families, which typically include both strongly cold-inducible members and members that are constitutively expressed or respond to other signals (39, 48). The differing induction profiles of RbpA, RbpB, and RbpC suggest a similar functional diversification within the *Bacteroides* RBP family, in which individual members may be tuned to distinct regulatory inputs.

A central unresolved question is the molecular mechanism by which RRM-1 RBPs influence gene expression during cold stress. Two models are possible, and they are not mutually exclusive. In the first, RBPs function analogously to canonical CSPs as RNA chaperones that globally remodel RNA secondary structure to maintain translational efficiency at low temperature, where thermodynamic stabilization of mRNA structures would otherwise impede ribosome progression (30). In the second model, RRM-1 RBPs function like the global post-transcriptional regulators Hfq (63), ProQ (64), and CsrA (65) which modulate gene expression by binding mRNAs to alter their stability or translational accessibility, and by facilitating or antagonizing the activity of small regulatory RNAs. The finding by Rüttiger, et al., (24) that *B. thetaiotaomicron* RbpB directly associates with both mRNAs and sRNAs and modulates the activity of at least one sRNA through a molecular sponge mechanism, suggests that RBPs may operate through sRNA-mediated regulatory pathways in addition to, or instead of, direct RNA chaperone activity. Our transcriptome data reveal extensive changes in gene expression upon loss of RBPs under cold conditions, but these changes almost certainly reflect a mixture of direct effects (genes whose expression is regulated through direct RBP-RNA interactions) and indirect effects arising from downstream regulatory consequences. The limited overlap between our differentially expressed genes and the published RbpB CLIP-seq dataset (24), which captures only RbpB targets at 37°C, likely reflects both the temperature-dependence of RBP-RNA interactions and the uncharacterized contributions of RbpA and RbpC. Distinguishing direct from indirect regulatory effects will require defining the RBP-RNA interactome under cold conditions for all three RBPs, for example, through CLIP-seq at low temperature, and will provide the mechanistic foundation needed to determine whether *Bacteroides* RRM-1 RBPs function primarily as cold shock-type RNA chaperones, as global post-transcriptional regulators analogous to Hfq and ProQ, or as proteins that integrate both functions.

The survival data connect this molecular story to an ecologically relevant context. Increasing evidence suggests that the human gut microbiome is shaped not only by vertical inheritance but also by horizontal transmission between adults. Several metagenomic studies have shown that adult cohabitation leads to detectable microbiome transmission, with *Bacteroidales* species displaying the highest transmissibility across cohabiting individuals (16, 17). Our finding that strains lacking RBPs show significantly reduced survival under combined cold and oxygen stress, while Δ*BT1884* strain does not, suggests that RBPs are the primary mediators of environmental resilience under conditions that *Bacteroides* encounter during host-to-host transmission. In contrast, *BT1884* appears to play a more limited role on its own, although its contribution becomes apparent when RBP function is absent.

Taken together, this work identifies a non-canonical cold stress regulatory system in *Bacteroides* built from RRM-1 domain proteins that are structurally and evolutionarily distinct from canonical CSPs yet recapitulate many of their key regulatory features. The functional redundancy within the RBP family, their coordinated relationship with BT1884, and their apparent role in cold tolerance and environmental resilience collectively suggest that post-transcriptional regulation by RRM-1 RBPs is a critical and underappreciated axis of *Bacteroides* physiology. Key questions remain: whether *rbp* 5’ UTRs function as RNA thermosensors, as suggested by the rapid accumulation of RBP proteins upon cold shock despite high transcript levels at 37 °C, analogous to the *cspA* 5’ UTR (60, 66); whether RBPs act as RNA chaperones or post-transcriptional regulators or both; and how their regulatory targets and mechanisms shift between optimal and cold growth conditions. Answering these questions will not only clarify how *Bacteroides* achieves cold adaptation in the absence of canonical machinery but will also shed light on the broader principles by which abundant and ecologically successful gut bacteria maintain regulatory flexibility under the diverse environmental challenges they encounter.

## MATERIALS AND METHODS

### Media and growth conditions

All strains, plasmids, and oligonucleotides used in this study are listed in Tables S7, S8, and S9, respectively. *B. thetaiotaomicron* VPI-5482 and other *Bacteroides* species were routinely cultured anaerobically at 37°C without shaking in a Coy Laboratory Products vinyl anaerobic chamber with an input gas mixture of 20% CO₂, 10% H₂, and 70% N₂, unless otherwise specified. Routine culturing of *Bacteroides* was performed in TYG broth, composed of tryptone, yeast extract, and glucose (67), and on brain heart infusion agar plates (BHI; Difco) supplemented with either 10% defibrinated horse blood (Quad Five) or 5 µg/mL hemin and 2.5 µg/mL vitamin K₃ (BHI-S). Plates were incubated anaerobically at 37°C. *Escherichia coli* strains were grown aerobically on BHI-S agar for conjugations and in Luria broth for all other applications. When needed, antibiotics or inducers were added at the following final concentrations: 100 µg/mL ampicillin (GoldBio), 200 µg/mL gentamicin (GoldBio), 25 µg/mL erythromycin (VWR), and 100 ng/mL anhydrotetracycline (Sigma).

### Construction of mutant strains and plasmids

Markerless chromosomal modifications were generated using the counterselectable allelic exchange vector pLGB13 (68). For deletion or insertion of SPOT epitope tag sequences, approximately 1 kb upstream and downstream flanking regions of the target gene were amplified by PCR and cloned into restriction-digested pLGB13 using the NEBuilder HiFi DNA Assembly Kit (NEB). Sequence-verified vectors were introduced into *Bacteroides* species via conjugation with *Escherichia coli* S17 λpir as previously described (28). Transconjugant merodiploids were selected on BHI-S agar plates containing gentamicin and erythromycin. Integration of the vector at the target locus was verified by colony PCR. For counterselection, a merodiploid colony harboring the appropriate vector was grown overnight in TYG broth, diluted 1:100 into 5 mL fresh TYG, and grown to exponential phase (OD600 ≈ 1). Subsequently, 3–5 µL of culture was plated onto BHI-S agar plates supplemented with 100 ng/mL anhydrotetracycline (Sigma) and incubated anaerobically for 2 days. Colonies carrying the desired chromosomal modifications were verified by colony PCR and sequencing. Complementation strains were generated using pNBU2-ErmG vectors (69) carrying the complementing genes, as previously described (28). Conjugated *Bacteroides* strains were selected on BHI-S agar plates containing gentamicin and erythromycin. Integration of the pNBU2 vector into the chromosomal attachment site was confirmed by colony PCR. Q5 High-Fidelity DNA Polymerase (NEB) was used for PCR reactions involved in cloning, whereas GoTaq® DNA Polymerase (Promega) was used for colony PCR.

### Luciferase assays

NanoLuc luciferase assays were performed as previously described (70). Strains were grown overnight in TYG medium. The following day, cultures were diluted 1:100 into 5 mL of fresh TYG and grown to exponential phase (OD600 = 0.6–1.0). A 100 µL aliquot of culture was centrifuged at 21,000 × g for 3 min, resuspended in 100 µL BugBuster Protein Extraction Reagent (Novagen), and lysed by a single freeze-thaw cycle. Lysates were nutated at room temperature for 10 min and clarified by centrifugation at 21,000 × g for 10 min at 4°C. NanoLuc® luciferase activity was measured using the Nano-Glo® Luciferase Assay System (Promega). Briefly, 10 µL of clarified lysate was mixed with an equal volume of Nano-Glo reagent in a 96-well microplate (Costar) and incubated for 5 min at 25°C prior to luminescence measurement. Luminescence was measured using a plate reader (BioTek Cytation 1 Cell imaging reader, Agilent) with an integration time of 1 s and a gain setting of 100. Relative luciferase activity was normalized to OD600. Data were analyzed using Microsoft Excel and GraphPad Prism.

### Ribosome extraction and sucrose sedimentation

Ribosome extraction and sucrose sedimentation were adapted from (52). Wild-type *Bacteroides thetaiotaomicron* VPI-5482 and the Δ*rbpABC* mutant were grown in TYG medium at either 37°C or 21°C. Exponential-phase samples were harvested at an OD600 of 0.6–1.0. Stationary-phase samples were harvested after 24 h of growth at 37°C or after 48 h of growth at 21°C.Equivalent culture volumes were harvested and resuspended in lysis buffer containing 50 mM Tris-HCl (pH 7.5), 10 mM MgCl₂, 100 mM NH₄Cl, and 6 mM β-mercaptoethanol. Cells were lysed by bead beating, and cell debris was removed by centrifugation at 21,000 × g for 30 min at 4°C to obtain clarified lysates. Clarified lysates were layered onto a 36% sucrose cushion and centrifuged at 120,000 × g for 16 h at 4°C using a Beckman 70Ti rotor. Ribosome pellets were washed twice with lysis buffer and resuspended by gentle rocking at 4°C for 1 h. Purified ribosomes were gently loaded onto 13 mL 15–50% (w/v) sucrose gradients prepared in lysis buffer and centrifuged at 71,000 × g for 16 h at 4°C using a Beckman SW41 rotor. Ribosome profiles were monitored by measuring absorbance at 260 nm using an ÄKTA purification system (GE Healthcare).

### Western blot analysis

Western blot analysis was performed to detect SPOT-tagged proteins. Strains carrying chromosomally SPOT-tagged alleles were grown anaerobically in TYG medium under the indicated conditions. Cells were harvested from equivalent culture volumes, pelleted by centrifugation, and resuspended in BugBuster Protein Extraction Reagent (Novagen) supplemented with EDTA-free protease inhibitor cocktail. Cells were lysed by bead beating, and lysates were clarified twice by centrifugation at 21,000 × g for 10 min at 4°C. Protein concentrations were determined using the Bradford assay, and 100 µg of total protein was mixed with SDS sample buffer and boiled for 10 min. Samples were separated by SDS-PAGE using 10–20% Tricine gels and transferred to membranes for immunoblotting. Membranes were blocked in milk-based blocking buffer and incubated with primary antibody against the SPOT tag, followed by horseradish peroxidase-conjugated secondary antibody. Signals were detected using SuperSignal West Pico PLUS chemiluminescent substrate (Thermo Fisher Scientific) and imaged using an iBright CL1000 imaging system.

### Cold induction assays

For cold induction assays, strains carrying chromosomally SPOT-tagged alleles were grown anaerobically overnight in TYG medium at 37°C. The following day, cultures were diluted 1:100 into fresh TYG medium and grown anaerobically at 37°C to exponential phase. Cultures were then transferred to either 21°C or 10°C, and samples were collected at the indicated time points after the temperature shift. Where indicated, control cultures were maintained at 37°C or grown continuously at the corresponding cold temperature. At each time point, aliquots were collected, cells were harvested by centrifugation, and whole-cell lysates were prepared as described above for Western blot analysis. Equal amounts of total protein were loaded for each sample, and SPOT-tagged protein levels were analyzed by Western blotting.

### Cold temperature viability assays

Viability assays were performed with three independent biological replicates. Wild-type *Bacteroides thetaiotaomicron* VPI-5482, Δ*rbpABC*, Δ*BT1884*, and Δ*rbpABC* Δ*BT1884* strains were grown anaerobically overnight in TYG broth at 37°C for 16–18 h. The following day, cultures were diluted 1:100 into fresh TYG broth and incubated for an additional 24 h at 37°C to stationary phase. Cultures were then divided into three aliquots and transferred to anaerobic incubation at 37°C, anaerobic incubation at 21°C, or oxygen-exposed incubation at 21°C. For oxygen exposure, cultures were incubated with shaking at 250 rpm. Prior to transfer, aliquots were collected to determine baseline viable cell counts. Cultures were maintained under the indicated conditions for 20 days, and aliquots were collected on days 6, 12, and 18 for CFU enumeration. Samples were serially diluted in TYG broth, and 100 µL aliquots of appropriate dilutions were plated on BHI-S agar. Plates were incubated anaerobically at 37°C for 2 days, and CFU/mL was calculated based on colony counts.

### RNA extraction, RNA sequencing sample preparation, processing and analysis

Strains were grown in three biological replicates overnight in TYG medium at 37°C. The following day, cultures were diluted 1:100 into 10 mL of fresh TYG and incubated at either 37°C or 21°C to exponential phase (OD600 = 0.6–1.0). Total RNA was isolated using the QIAGEN RNeasy Mini Kit (catalog no. 74104) with on-column DNase I digestion. RNA was quantified using a Qubit 2.0 fluorometer (Invitrogen) and submitted to the Roy J. Carver Biotechnology Center at the University of Illinois Urbana-Champaign for library preparation and sequencing. Ribosomal RNA was removed using the FastSelect Bacteria kit (QIAGEN). RNA-seq libraries were prepared using the KAPA Hyper Stranded mRNA Library Kit (Roche) without mRNA capture. Libraries were pooled, quantified by qPCR, and sequenced on a partial 10B lane for 101 cycles from one end of the fragments on a NovaSeq X Plus using V1.0 sequencing kits. FASTQ files were generated and demultiplexed using bcl2fastq Conversion Software v2.20 (Illumina). Reads were filtered and trimmed using fastp v0.23.4 with default settings and aligned to the *B. thetaiotaomicron* reference genome GCF_000011065.1 using Bowtie2 v2.5.5 with default settings. Aligned reads were quantified using featureCounts v2.0.3 with the -s parameter set to 2. Differential expression analysis was performed in R v4.3.2 using DESeq2 v1.44.0. Genes were considered differentially expressed if they had an absolute log2 fold change > 1 and an adjusted P value < 0.05. Over-representation analysis (ORA) of differentially expressed genes was performed using clusterProfiler v4.20 with gene sets from the Gene Ontology (GO) database.

Similarly, gene set enrichment analysis (GSEA) was performed using clusterProfiler and a custom script on a ranked list of all *B. thetaiotaomicron* genes based on log2 fold change, using gene sets from the GO database.

### Statistical methods and figures

Statistical analyses were performed using GraphPad Prism v10.2.2. The statistical tests used for each experiment are indicated in the corresponding figure legends. Graphs were generated using GraphPad Prism. Figures were assembled using Inkscape, Microsoft PowerPoint, and BioRender. Claude was used to assist with figure layout and schematic formatting. All AI-assisted content was reviewed and edited by the authors.

## ACKNOWLEDGEMENTS

We thank the members of HL’s thesis committee, Bill Metcalf, James Slauch, and Paola Mera for their advice and guidance throughout the course of this project. We are grateful to Patrick Degnan at the University of California, Riverside, for generously sharing bacterial strains. We also thank Alvaro Hernandez, Chris Wright, and the Roy J. Carver Biotechnology Center staff for their support with RNA-seq experiments. We thank Raven Huang at the University of Illinois Urbana-Champaign for helpful guidance and access to equipment for ribosome sedimentation experiments. We thank James Imlay at the University of Illinois Urbana-Champaign for his mentorship and guidance. We also thank former and current members of the Vanderpool laboratory for helpful discussions and support.

## Data availability

RNA sequencing data generated in this study are publicly available in the NCBI Sequence Read Archive under BioProject accession number PRJNA1480845.

## Supplemental Material

